# *De novo* identification of satellite DNAs in the sequenced genomes of *Drosophila virilis* and *D. americana* using the RepeatExplorer and TAREAN pipelines

**DOI:** 10.1101/781146

**Authors:** Bráulio S.M.L. Silva, Pedro Heringer, Guilherme B. Dias, Marta Svartman, Gustavo C.S. Kuhn

## Abstract

Satellite DNAs are among the most abundant repetitive DNAs found in eukaryote genomes, where they participate in a variety of biological roles, from being components of important chromosome structures to gene regulation. Experimental methodologies used before the genomic era were not sufficient despite being too laborious and time-consuming to recover the collection of all satDNAs from a genome. Today, the availability of whole sequenced genomes combined with the development of specific bioinformatic tools are expected to foster the identification of virtually all of the “satellitome” from a particular species. While whole genome assemblies are important to obtain a global view of genome organization, most assemblies are incomplete and lack repetitive regions. Here, we applied short-read sequencing and similarity clustering in order to perform a *de novo* identification of the most abundant satellite families in two *Drosophila* species from the *virilis* group: *Drosophila virilis* and *D. americana*. These species were chosen because they have been used as a model to understand satDNA biology since early 70’s. We combined computational tandem repeat detection via similarity-based read clustering (implemented in Tandem Repeat Analyzer pipeline – “TAREAN”) with data from the literature and chromosome mapping to obtain an overview of satDNAs in *D. virilis* and *D. americana*. The fact that all of the abundant tandem repeats we detected were previously identified in the literature allowed us to evaluate the efficiency of TAREAN in correctly identifying true satDNAs. Our results indicate that raw sequencing reads can be efficiently used to detect satDNAs, but that abundant tandem repeats present in dispersed arrays or associated with transposable elements are frequent false positives. We demonstrate that TAREAN with its parent method RepeatExplorer, may be used as resources to detect tandem repeats associated with transposable elements and also to reveal families of dispersed tandem repeats.

## Introduction

The genome of eukaryotes encloses a variety of repetitive DNA sequences which comprises most of the nuclear DNA of several organisms, including animals, plants and insects [1,2]. Among them are the satellite DNAs (satDNAs), usually defined as abundant, tandemly repeated noncoding DNA sequences, forming large arrays (hundreds of kilobases up to megabases), typically located in the heterochromatic regions of the chromosomes [3,4], although short arrays may additionally be present in the euchromatin [5,6].

The collection of satDNAs in the genome, also known as the “satellitome”, usually represents a significant fraction (>30%) of several animal and plant genomes. Other classes of noncoding tandem repeats include the microsatellites, with repeat units less than 10 bp long, array sizes around 100 bp and scattered distributed throughout the genome; and the minisatellites, with repeats between 10 to 100 bp long, forming up to kb-size arrays, located at several euchromatic regions, with a high density at terminal chromosome regions [3,4]. Therefore, the best criteria to distinguish satellites from micro and minisatellites are long array sizes and preferential accumulation at heterochromatin for the former.

SatDNAs do not encode proteins, but they may play important functional roles in the chromosomes, most notably related to chromatin modulation and the establishment of centromeres [7–9]. They are among the fastest evolving components of the genome (although some conserved satellites have also been reported) [10–12], and such behavior combined to their abundance and structural role have major implications for the evolution and diversification of genomes and species [8,13].

Since the discovery of satDNAs in the early 60’s, species from the genus *Drosophila* have been used as a model to address several aspects of satDNA biology, such as their origin, organization, variation, evolution and function [e.g. 7,14–18].

Today, several *Drosophila* genomes have been sequenced by next-generation technologies and new bioinformatic tools have been designed for the identification of repetitive DNAs from this vast source of genomic resources. Among them, the RepeatExplorer software [19] has been successfully used for *de novo* identification of repetitive DNAs directly from unassembled short sequence reads, and the recently implemented TAREAN pipeline [20] was introduced to specifically identify putative satDNAs. Such a combination between sequenced genomes and bioinformatic tools is now expected to foster the identification of the full “satellitome” of any given species [e.g. 21–25]. Despite all such facilities in hand, only a few *Drosophila* species had their satDNA landscape determined using these new approaches [22].

In the genus *Drosophila*, genome sizes vary between ~130 Mb to ~400 Mb, but most analyzed species have genome sizes around 180-200 Mb, such as *D. melanogaster* [26,27]. The satDNA content also varies across species, from ~2% in *D. buzzatii* [22] to ~60% in *D. nasutoides* [28]. Some studies suggest a positive correlation between genome size and the amount of satDNAs in *Drosophila* [27,29,30].

The genome size of *D. virilis* (*virilis* group), with ~400 Mb, is among the largest reported for *Drosophila*. Accordingly, the estimated satDNA in this species is also high (>40%) [27,31]. Previous studies using CsCl density gradients revealed that three evolutionary related satDNAs with repeat units 7 bp long and only one mutation difference, named satellite1 (5’ ACAAACT 3’), satellite2 (5’ ATAAACT 3’) and satellite3 (5’ ACAAATT 3’) together represent ~40% of its genome [31,32]. These satellites mapped predominantly to the heterochromatic regions of all chromosomes except the Y. Another satDNA identified in this species, but using genomic DNA digestion with restriction endonucleases, was named pvB370, and consists of 370 bp long repeat units [33] predominantly located at sub-telomeric regions and, to a lesser extent, along some discrete euchromatic loci [34]. Other abundant tandem repeats (TRs) have been identified in the *D. virilis* genome, such as the 220TR and 154TR families, which belong to the internal structure of transposable elements [16,35], the 225 bp family, present in the intergenic spacer of ribosomal genes, and the less characterized 172 bp family [36]. A recent study reported additional tandem repeats less than 10 bp long but at low abundance [18].

The high throughput and low cost of current whole-genome sequencing technologies have made it possible to obtain genome assemblies for a wide range of organisms. However, *de novo* whole-genome shotgun strategies are still largely unable to fully recover highly repetitive regions such as centromeres and peri-centromeric regions and, as a result, satDNAs are usually misrepresented or absent from such assemblies [37]. One way of circumventing the assembly bottleneck is to directly identify repeats from raw sequencing reads. One of such approaches is implemented in the RepeatExplorer pipeline, already used in a wide range of plant and animal species [21,38,39]. RepeatExplorer performs similarity-based clustering of raw short sequencing reads and partial consensus assembly, allowing for repeat identification even from small samples of genome coverage. A recent development of RepeatExplorer includes the TAREAN pipeline for the specific detection of tandem repeats by searching for circular structures in directed read clusters [20].

In the present study, we aim to test the ability of TAREAN to correctly identify satDNAs in *D. virilis*. To refine and expand our knowledge of the identified putative satDNAs, in some cases we mapped them into the mitotic and polytene chromosomes using the FISH (Fluorescent in situ Hybridization) technique.

There are several examples showing that satDNA abundance may vary widely even across closely related species [10,40]. For example, one species might present few repeats in the genome (therefore not being identified as a satellite), while a close related species present thousands, reaching a satDNA status. For this reason, we also add to our study *D. americana*, a species belonging from the *virilis* group, but separated from *D. virilis* by ~4.1 Mya [41].

## Material and Methods

### RepeatExplorer and Tandem Repeat Analyzer (TAREAN) analyses

The *in silico* identification of putative satDNAs was performed using the RepeatExplorer and Tandem Repeat Analyzer (TAREAN) pipelines [19,20] implemented in the Galaxy platform [42]. These algorithms were developed to identify and characterize repetitive DNA elements from unassembled short read sequences. We used the publicly available *Drosophila virilis* strain 160 (SRX669289) and *Drosophila americana* strain H5 (ERX1035147) Illumina paired-end sequences. The sequences were obtained through the “European Nucleotide Archive” (EBI) database and their quality scores measured with the “FASTQC” tool. We used “FASTQ Groomer” (Sanger & Illumina 1.8 +) to convert all the sequences to a single fastqsanger format. We removed adapters and excluded any reads with more than 5% of its sequence in low quality bases (Phred cutoff < 10) using the “Preprocessing of fastq paired-reads” tool included in the RepeatExplorer Galaxy instance. The interlaced filtered paired-end reads were used as input data for the RepeatExplorer clustering and Tandem Repeat Analyzer tools with the following settings: “sample size = 2,000,000 – select taxon and protein domain database version (REXdb): Metazoa version 3.0 – select queue: extra-long and slow”. For the TAREAN analyses we also used the “perform cluster merging” tool for reducing the redundancy of the results.

The results were provided in a HTML archive report and all the data was downloaded in a single archive for further investigation. Here, we analyzed clusters with abundances representing >0.5% of the genome of *Drosophila virilis*.

Clusters with tandem repeats identified by TAREAN are denoted as putative high or low confidence satellites. These estimates are denoted according to the “Connected component index (C)” and “Pair completeness index (P)”. The C index indicates clusters formed by tandemly repeated genomic sequences, while the P index measures the ratio between complete read pairs in the cluster and the number of broken pairs, that is directly related to the length of continuous tandem arrays [20].

### Molecular Techniques

We extracted total genomic DNA from a pool of 20 adult flies of *D. virilis* (strain 15010-1051.51 from Santiago, Chile) and *Drosophila americana* (strain H5 from Mississipi, United States of America) with the Wizard® Genomic DNA Purification Kit (Promega Corporation). We used the consensus sequences from each satDNA identified by RepeatExplorer/TAREAN for primer’s design. Satellite DNAs were PCR amplified with the following primers: forward (F) and reverse(R): Sat1_F(ACAAACTACAAACTACAAACTACAAACTACAAACT),Sat1_R(AGTTTGTAGTTTGTAGTTTGTAGTTTGTAGTTTGT),172TR_F(ATTTATGGGCTGGGAAGCTTTGACGTATG),172TR_R(CGGTCAAATCTCATCCGATTTTCATGAGG),225TR_F(GCGACACCACTCCCTATATAGG),225TR_R(CGCGCAAGGCATG TCATATG).

The PCR products were excised from agarose gels and ligated into pGEM-T vector plasmids (Promega) with T4 DNA ligase (Promega). For cloning, the plasmids were multiplied into *E.coli* cells and then eluted with the PureLink™ Quick Plasmid Miniprep Kit (Invitrogen). To ensure the presence of the inserts, the final samples were Sanger sequenced in an ABI3130 and later analyzed in the Chromas software (Technelysium). Clones with satDNAs inserts were later prepared as probes for FISH experiments.

### Fluorescent in situ hybridization (FISH) experiments

The metaphase and polytene chromosomes were obtained from neuroblasts and salivary glands of third instar larvae [43,44] from *D.virilis* (strain 15010-1051.51) and *D. americana* (strain H5). The labeling of probes and FISH experiment conditions were conducted according to [16]. The satDNA probes were immunodetected with antidigoxigenin-Rhodamine and avidin-FITC (Roche Applied Science).

We used DAPI “40,6-diamidino-2-phenylindole” (Roche) with “SlowFade” antifade reagent (Invitrogen) for DNA counterstaining. The slides analyses were made with an Axio Imager A2 epifluorescence microscope equipped with the AxiocamMRm camera (Zeiss). Images were captured with Axiovision (Zeiss) and edited in Adobe Photoshop.

## Results

### Identification of putative satDNAs in *D. virilis* and *D. americana*

The most abundant putative satDNAs (covering >0.5% of the genome) identified by the TAREAN pipeline are shown in Table 1 (see S1 Fig for histogram summary analyses and S4-S15 Figs for detailed data from each cluster retrieved). All of the six identified tandem repeat families (Sat1, 154TR, pvB370, 172TR, 225TR, 36TR) are common to the two species and have been previously identified.

**Table 1.** Putative satellite DNAs in *D. virilis* and *D. americana* identified by TAREAN.

|  | *Drosophila virilis* |  |  |  |  |  | *Drosophila americana* |  |  |  |  |  |
| --- | --- | --- | --- | --- | --- | --- | --- | --- | --- | --- | --- | --- |
| Tandem repeat family<sup>a</sup> | Sat1 | 154TR | pvB370 | 172TR | 225TR | 36TR | Sat1 | 172TR | 154TR | pvB370 | 225TR | 36TR |
| Satellite confidence | High | Low | Low | High | Low | Low* | High | Low | Low | High | High* | n/a* |
| Satellite probability | 0.92 | 0.03 | 0.53 | 0.73 | 0.69 | 0.00* | 0.91 | 0.69 | 0.04 | 0.75 | 0.76* | 0.00* |
| C index<sup>b</sup> | 0.98 | 0.94 | 0.96 | 0.97 | 0.99 | 0.94* | 0.96 | 0.97 | 0.94 | 0.97 | 0.97* | 0.72* |
| P index<sup>b</sup> | 0.92 | 0.71 | 0.81 | 0.86 | 0.87 | 0.52* | 0.97 | 0.85 | 0.72 | 0.87 | 0.86* | 0.24* |
| Consensus size | 7bp | 154bp | 370bp | 171bp | 225bp | 36bp* | 7bp | 171bp | 154bp | 199bp | 225bp* | n/a* |
| Genome proportion (%) | 12.0 | 1.6 | 1.6 | 1.1 | 0.8 | 0.7* | 9.0 | 2.7 | 2.2 | 1.7 | 0.9* | 0.4* |

\*. Results obtained from RepeatExplorer instead of TAREAN.
<sup>a</sup>. Ordered by abundance from higher to lower.
<sup>b</sup>. C and P indexes are explained in Materials and Methods.

Although the total abundance of these tandem repeats is similar (~17%) in the two species, there are differences in the estimated proportion occupied by each putative satDNAs between the species.

To further characterize the tandem repeat families identified *in silico*, we constructed DNA probes using the consensus sequences generated by TAREAN from three families and used them to verify their localization in metaphase and polytene chromosomes. In the following sections we describe our *in silico* and FISH analyses for each identified family, comparing the results with previous studies and discussing if TAREAN correctly identified and distinguished satDNAs from other classes of tandem repeats. The tandem repeat families are described below in order of their abundance (higher to lower) as revealed for *D. virilis*.

### Sat1

The most abundant tandem repeat identified by TAREAN in *D. virilis* and *D. americana* is composed by a 7 bp long repeat corresponding to the previous described satellite I [32]. In *D. virilis*, our FISH experiments in metaphase chromosomes showed this satDNA occupying the pericentromeric region of all autosomes except the small dot chromosomes, and in the X and Y chromosomes (Fig 1A). However, the hybridization in polytene chromosomes revealed that Sat1 also localizes in the pericentromeric region of the dot chromosome (Fig 2A). [31] showed a similar hybridization pattern, although their results did not consistently demonstrate Sat1 signals in the dot and Y chromosomes.

**Fig 1.**
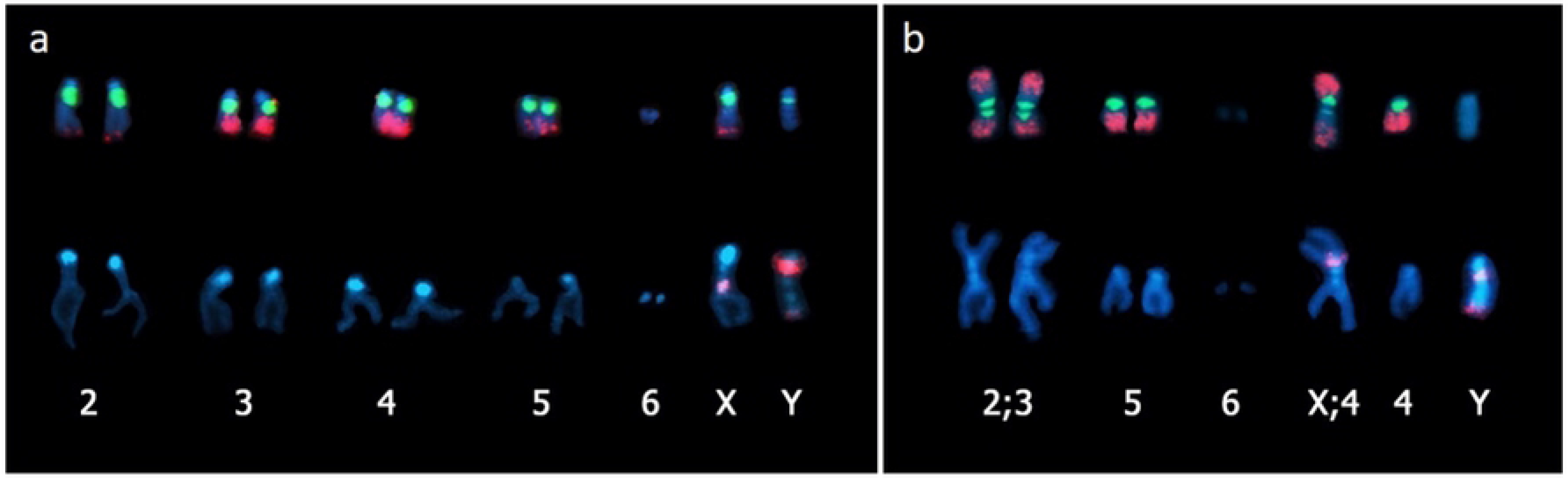
Chromosome location of Sat1, 172TR and 225TR by FISH on metaphase chromosomes. (a) *Drosophila virilis* and (b) *Drosophila americana*. Upper panel: Sat 1 (green) and 172TR (red). Lower panel: 225TR (red).

**Fig 2.**
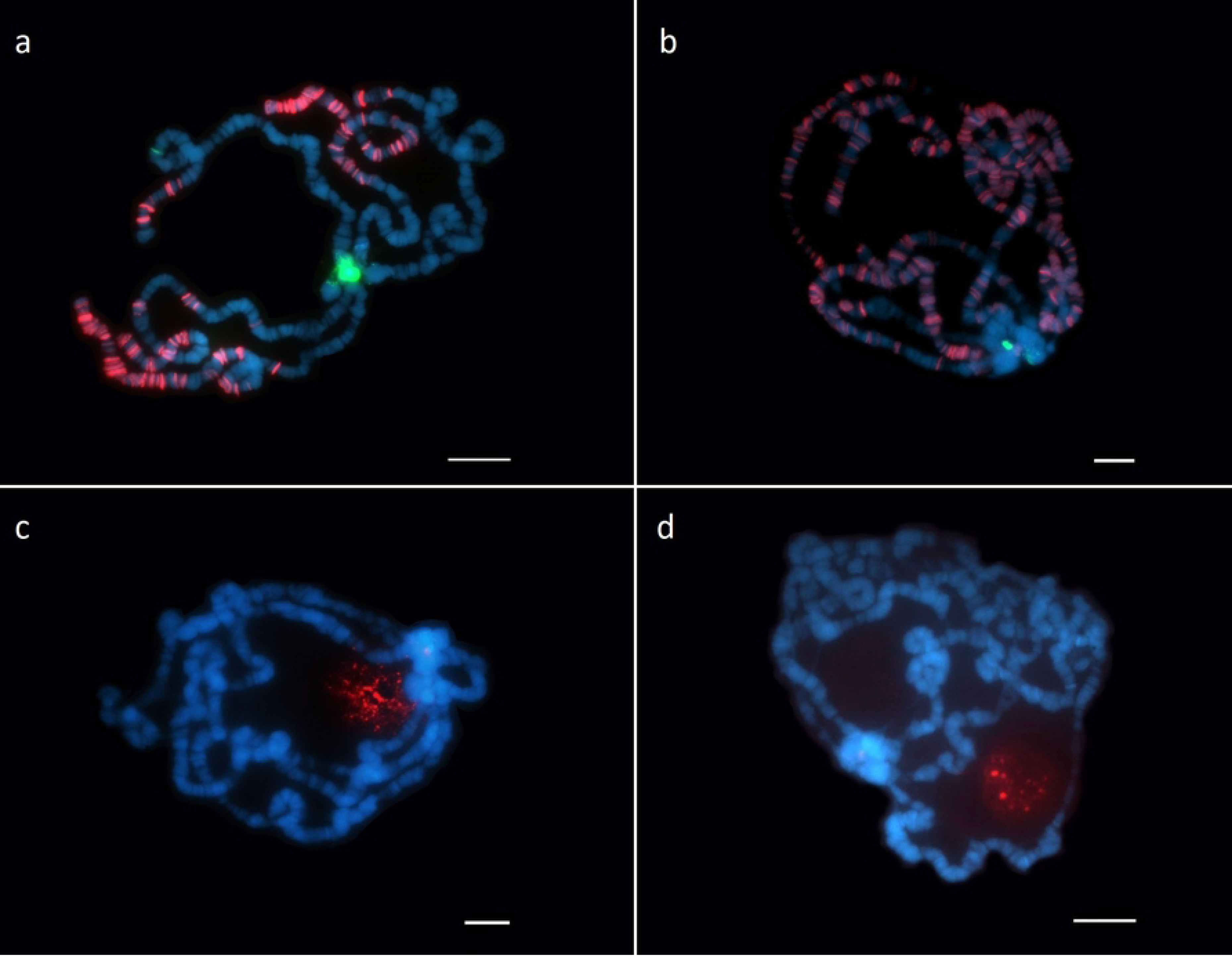
Chromosome location of Sat1, 172TR and 225TR by FISH on polytene chromosomes. (a,c) *Drosophila virilis* and (b,d) *Drosophila americana*. (a-b) Sat 1 (green) and 172TR (red). (c-d) 225TR (red). Scale bars represent 10μm.

In *D. americana*, Sat1 signals were detected in the pericentromeric region of all autosomes in metaphase chromosomes, except the dot (Fig 1B), while in polytene chromosomes, Sat1 signals were also observed in the dot chromosomes (Fig 2B). However, differently to what was observed in *D. virilis*, our Sat1 hybridizations in the *D. americana* polytene dot chromosomes did not give enough information about the precise location of this satDNA, although it also appears to occupy a portion of the pericentromeric region. As another difference from *D. virilis*, Sat1 sequences appear to be absent from the Y chromosome in *D. americana* (Fig 1B). Our FISH results corroborate the smaller genomic fraction occupied by this satDNA in *D. americana* (~9% against ~12% in *D. virilis*), revealed by the *in silico* analysis (Fig 1, 2A and 2B). These new findings in *D. americana* and *D. virilis* also agree with recent results from [45].

### 154TR

The genomic distribution of 154TR has been already recently studied in detail in *D. virilis* and *D. americana* using FISH in metaphase and polytene chromosomes [35]. This sequence was independently identified *in silico* by [36] and [46]. The 154TR was characterized as a tandem repeat derived from a Helitron transposable element [36], which was studied in detail and classified as a family named DINE-TR1 [35]. DINE-TR1 elements containing 154TR homologous sequences were found in several Acalyptratae species, mostly within the *Drosophila* genus, although long arrays (> 10 copies) of 154TR were only detected in three species (*D. virilis*, *D. americana* and *D. biarmipes*) [35]. FISH in metaphase and polytene chromosomes revealed that 154TR is located in the distal pericentromeric region (β-heterochromatin) and many euchromatic loci of all autosomes and the X chromosome of *D. virilis* and *D. americana*. In addition, this tandem repeat covers a large portion of the Y chromosome in both species. In *D. virilis*, 154TR signals are very abundant in the centromeric heterochromatin of chromosome 5 and are also found in a discrete region within the pericentromeric region of the X chromosome [35].

Our results from the TAREAN analysis classified 154TR as a putative satellite with low confidence in both species (Table 1). We suggest that this result is probably a consequence of 154TR being both tandemly repeated, like a satDNA, and dispersed, like a transposable element. In this case, even though the connected component index (*C*) of 154TR is high, its relatively low pair completeness index (P) contributes to its classification as a putative satellite with low confidence by TAREAN (Table 1). We suggest that 154TR is not a satDNA and thus, should be classified as a highly abundant dispersed tandem repeat.

### pvB370

The pvB370 satellite was first described by [33], which also identified this family as deriving from the direct terminal repeats of pDv transposable elements [47]. In a following study, [34] showed that in *D. virilis* and *D. americana*, pvB370 is located at several euchromatic loci and at the telomeric region of all chromosomes. Our *in silico* analyses indicate that pvB370 covers a similar portion of *D. virilis* (~1.6%) and *D. americana* (~1.7%) genomes, which agrees with Southern blot results from [33] that indicated a similar number of copies of pvB370 in both species.

Because pvB370 was previously mapped in the chromosomes of *D. virilis* and *D. americana* using FISH, we did not conduct a throughout analysis on both species. However, because pvB370 seems to display a euchromatic distribution [34] similar to the one we observed for 172TR (Fig 2A and 2B) we hybridized both pvB370 and 172TR probes concomitantly in *D. americana* polytene chromosomes. Our results showed no or little overlap between pvB370 and 172TR, although many arrays from the two families are very close (at least a few kbp) to each other (Fig 3).

**Fig 3.**
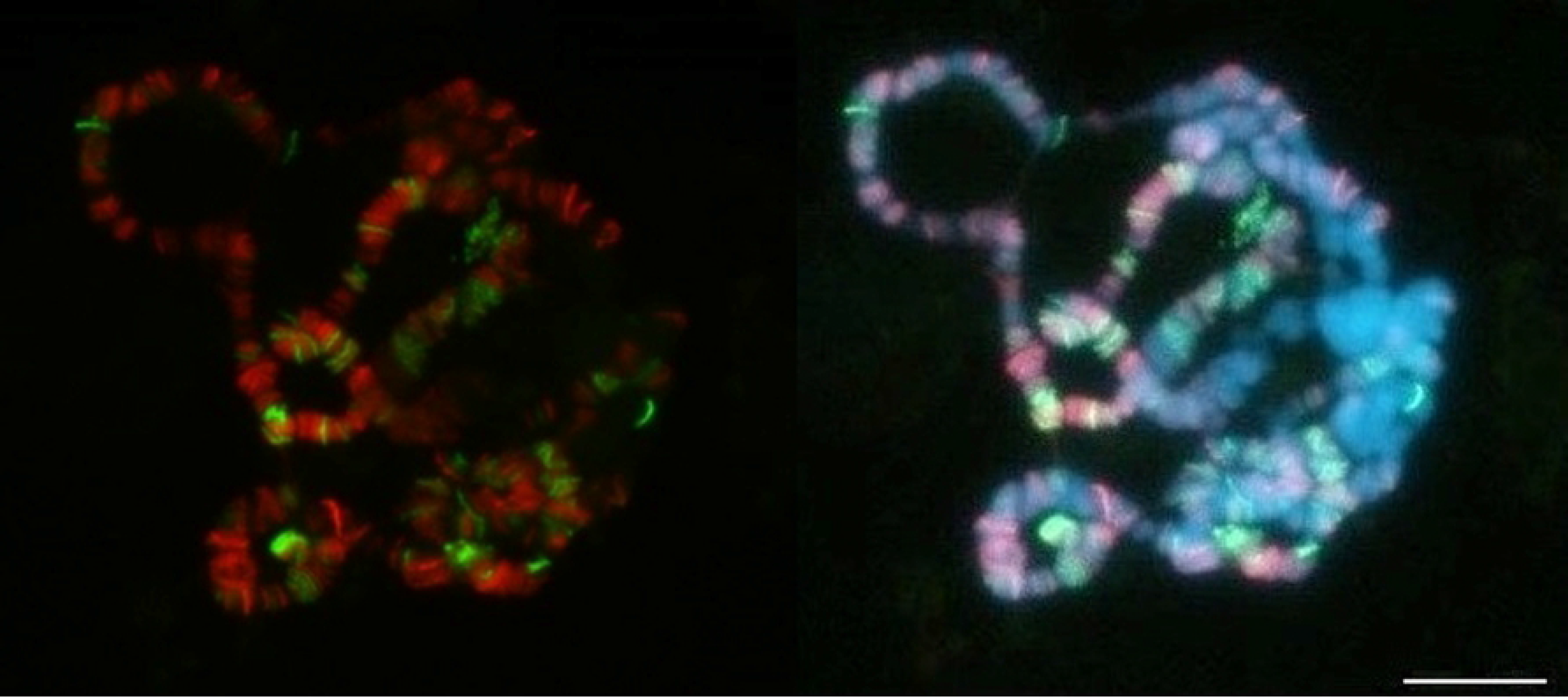
Chromosome location of 172TR and pvB370 by FISH on polytene chromosomes of *Drosophila americana*. There is no or little overlap between these tandem repeats. Red (172TR) and green (pvB370). Scale bar represents 10μm.

### 172TR

The 172TR family corresponds to the 172 bp tandem repeats previously identified *in silico* by [36]. Our FISH results in the metaphase and polytene chromosomes of *D. virilis* revealed that 172TR is distributed throughout the arms of autosomes 3, 4 and 5, in several loci at the X chromosome and at least in two loci in chromosome 2, including the subtelomeric region (Fig 1A and 2A). Most of its arrays are located at distal chromosome regions. No hybridization signals were detected in the dot and Y chromosomes.

The FISH results in *D. americana* showed 172TR signals at multiple loci along all autosomes, except the dot, and more equally distributed in both distal and proximal regions of chromosome arms (Fig 1B and 2B). Similarly to *D. virilis*, no hybridization signal was detected in the Y chromosome (Fig 1B). The FISH data (Fig 1, 2A and 2B) clearly showed a higher number of 172TR loci in *D. americana* compared to *D. virilis*, a result that is consistent with the higher overall abundance of 172TR repeats in *D. americana* predicted by the *in silico* analysis (Table 1).

### 225TR

The putative satDNAs detected in our *in silico* analyses as 225TR was previously identified as a component of intergenic spacers (IGS) of ribosomal genes from *D. virilis* located at the chromocenter and nucleolus regions of polytene chromosomes [36]. Our FISH experiments in polytene chromosomes confirmed these results in *D. virilis* (Fig 2C), additionally showing that in *D. americana* this family displays the same pattern of localization (Fig 2D).

In addition, we also performed FISH with a 225TR probe in metaphase chromosomes of both species for the first time, what revealed its location in the pericentromeric region of the X chromosome and in the pericentromeric and telomeric regions of chromosome Y (Fig 1A and 1B). This result is in accordance with previous studies showing the location of these IGS sequences in the sex chromosomes in *Drosophila* [48].

Although the TAREAN pipeline failed to detect the 225TR in *D. americana*, RepeatExplorer revealed the presence of this family. This indicates a possible limitation of TAREAN in detecting less abundant tandem repeats in comparison with RepeatExplorer. Moreover, TAREAN only retrieves clusters with highly circular structures, and therefore excludes 225TR-like repetitions that are associated with linear sequences (S12 and S13 Figs).

BLAST searches using a 225TR consensus sequence as a query against the genome of *D. virilis* resulted in several contigs containing up to 40 tandem repeats (data not shown). However, most of the arrays are dispersed and containing up to 10 tandemly repeated copies of the 225TR sequence, which agrees with its classification as a putative satellite with low confidence. Together, these observations indicate that, although 225TR is an abundant tandem repeat, it does not have all the typical features of a satDNA.

### 36TR

A putative satDNA with 36 bp long repeat units was detected by TAREAN but with low confidence in *Drosophila virilis* and occupying ~0.73% of *D. virilis* and ~0.48% of *D. americana* genomes (Table 1). However, similarly to pvB370, these 36 bp repeats correspond to an internal portion of the pDv transposable element [33,47].

Interestingly, the RepeatExplorer pipeline revealed that the cluster corresponding to this 36 bp tandem repeat has a high number of shared reads with the pvB370 cluster (S2 and S3 Figs). This result illustrates that RepeatExplorer is able to detect putative relationships between distinct repetitive sequences. In this case, the link between 36TR and pvB370 clusters is explained by their co-occurrence of these complete (36bp) and partial (pvB370) sequences within the pDv transposable element [33].

## Discussion

Here we performed *de novo* identification of the most abundant tandem repeat families in *D. virilis* and *D. americana*. These species were chosen because they have larger genomes compared to other *Drosophila* species and because they have been used as a model to understand satDNA biology since early 70’s. In order to do that, we combined the TAREAN results with data from the literature and, in some cases, with new chromosome mapping data obtained by us using FISH in metaphase and polytene chromosomes.

Because all of the repeats identified here had been previously detected by other methods, we were able to test if the TAREAN pipeline could correctly classify them as satDNAs or not.

TAREAN identified the heptanucleotide Sat1 as a satDNA with high confidence, which agrees with all attributes known for this family and the satDNA definition (i.e. high copy-number, long-arrays, predominant heterochromatic location) [31,32]. The Sat1 was identified as the most abundant tandem repeat in both *D. virilis* and *D. americana*, which is also in accordance with previous work [31,32]. However, the other two less abundant heptanucleotide satellites, Sat2 and Sat3, were not detected by TAREAN. As these three satellites differ from each other by a single substitution, they were likely all included in the Sat1 cluster by TAREAN. It is also worth mentioning that the heptanucleotide satDNA genomic fraction revealed by TAREAN (~12% for *D. virilis* and ~9% for *D. americana*) are significantly below the previously estimated of >40% genomic fraction, based on density gradient ultracentrifugation methods [31,49]. Although TAREAN might not be ideally suitable to quantify satellites with short repeat units [20], it is worth mentioning that [45] have recently demonstrated that Illumina sequence reads containing the heptanucleotide satellites from *D. virilis* tend to be highly enriched for low quality scores. Furthermore, the use of raw reads from different sequencing platforms did not allowed the recovery of simple satellites at the predicted ~40% genomic fraction indicated by previous works [45]. The gap between these estimates (12% to 40%) might reflect an intrinsic bias in current sequencing methods. A second possibility, which does not reject the first, is the existence of real differences in satDNA content between the different *D. virilis* strains used in these studies.

TAREAN classified the 154TR, pvB370 and 36TR families as putative satellites in *D. virilis* and *D. americana*. With the exception of pvB370 in *D. americana*, which was classified with high confidence, all remaining repeats had low confidence calls from TAREAN (Table 1). These tandem repeats are known to be abundant and associated with transposable elements (as integral parts or evolutionarily related), suggesting that RepeatExplorer and TAREAN could be used as resources to detect tandem repeats associated with transposable elements. In the case of 154TR, pvB370, and 36TR, the relationship could be assigned directly in the RepeatExplorer pipeline by identifying clusters of tandem repeats sharing a high number of reads with clusters associated to transposable elements (see S2 and S3 Figs), or indirectly in the TAREAN pipeline, by investigating the tandem repeats classified as putative satellites with low confidence (or lower values of satellite probability). The rationale behind this last procedure is that identified families with a ‘low satellite score’ might represent repetitive DNAs with intermediate features, being both highly dispersed but tandemly repeated. One situation in which this scenario is expected is the case where tandem repeats belonging to the terminal or internal portions of transposable elements underwent array expansion [35,50]. Nonetheless, some highly dispersed tandem repeats are not necessarily associated with transposable elements, which is the case of 172TR shown here and the 1.688 satDNA from *D. melanogaster* [5].

It is interesting to note that, in *D. virilis* and *D. americana*, the families 172TR, pvB370 and 154TR were either classified as putative satellites with low confidence, or with high confidence but associated with a relatively low satellite probability (Table 1). Because all these three families were found scattered distributed along the euchromatic arms of chromosomes, we suggest that a low ‘satellite score’ in the TAREAN pipeline is a good predictor of dispersed tandem repeats. As mentioned above, although there is no indication of a relationship between the 172TR family with any known transposable element, its lower satellite score from the *in silico* analysis correctly predicts the dispersed array distribution.

In conclusion, six abundant putative satDNAs were identified in *D. virilis* and *D. americana* by TAREAN: Sat1, 154TR, pvB370, 172TR, 225TR and 36TR. All of them have been previously characterized to a higher or lesser extent in previous works, but using different methodologies. The main advantage of TAREAN in comparison with previous methods aiming to identify satDNAs in *D. virilis* refers to its relative lack of bias compared to the *in silico* digestion applied by [36], that identify only tandem repeats presenting restriction sites, and the k-Seek method applied by [18] that specifically identify short tandem repeats less than 20 bp.

While Sat1 (identified by TAREAN as a satDNA with high confidence) is in fact a family that matches all features typically attributed for satDNAs, the classification of the other families as satDNAs (identified as a satDNA with low confidence on at least one species) is more controversial. The 154TR, pvB370 and 36TR families are associated with the internal structure of TEs, thus being distributed along the chromosome arms at different degrees of dispersion. The 225TR belongs to the IGS of ribosomal genes. In contrast, the 172TR family is an abundant tandem repeat but with exclusive euchromatic location, where they apparently do not to reach satDNA-like long arrays. Based on the repeat unit length of 172TR (172 bp), this family cannot be considered as a micro or minisatellite. In this context, it would be interesting to further investigate these five families (154TR, pvB370, 172TR, 225TR and 36TR) using long-read sequencing technologies, since they are expected to provide more detailed information about their copy number and array sizes.

## Supporting information

**S1 Fig. TAREAN histogram summary analyses of (a) *Drosophila virilis* (strain 160) and (b) *Drosophila americana* (strain H5).**

**S2 Fig. pvB370 and 36TR supercluster analysis in *Drosophila virilis* strain 160.**

**S3 Fig. pvB370 and 36TR supercluster analysis in *Drosophila americana* strain H5.**

**S4 Fig. Sat1 cluster analysis in *Drosophila virilis* strain 160.**

**S5 Fig. Sat1 cluster analysis in *Drosophila americana* strain H5.**

**S6 Fig. 154TR cluster analysis in *Drosophila virilis* strain 160.**

**S7 Fig. 154TR cluster analysis in *Drosophila americana* strain H5.**

**S8 Fig. pvB370 cluster analysis in *Drosophila virilis* strain 160.**

**S9 Fig. pvB370 cluster analysis in *Drosophila americana* strain H5.**

**S10 Fig. 172TR cluster analysis in *Drosophila virilis* strain 160.**

**S11 Fig. 172TR cluster analysis in *Drosophila americana* strain H5.**

**S12 Fig. 225TR cluster analysis in *Drosophila virilis* strain 160.**

**S13 Fig. 225TR cluster analysis in *Drosophila americana* strain H5.**

**S14 Fig. 36TR cluster analysis in *Drosophila virilis* strain 160.**

**S15 Fig. 36TR cluster analysis in Drosophila americana strain H5.**

